# A DIY guide for image-based spatial transcriptomic: TLS as a case example

**DOI:** 10.1101/2024.07.03.601914

**Authors:** Thomas Defard, Auxence Desrentes, Charles Fouillade, Florian Mueller

## Abstract

Spatial RNA profiling methods provide insight to the cellular heterogeneity and spatial architecture of complex, multi-cellular systems. Combining molecular and spatial information provides important clues to study tissue architecture in development and disease. Here, we present a comprehensive do-it-yourself guide to perform such experiments at reduced costs leveraging open-source approaches. This guide spans the entire life cycle of a project, from its initial definition to experimental choices, wet lab approaches, instrumentation and analysis. As a concrete example, we focus on Tertiary lymphoid structures (TLS), which we use to develop typical questions that can be addressed by these approaches.

## Introduction

### Importance of spatial context in biology

Spatial organization at the cellular level is a fundamental aspect of complex life, orchestrating intricate biological processes with remarkable precision. This intricate organization allows cells to efficiently exchange nutrients and signals, crucial for the proper functioning of multicellular organisms. Being able to describe the spatial landscape of the cells is hence essential to obtain a better understanding of tissue function (***1***).

In the context of the immune system, the effector functions are primarily carried out by motile patrolling cells dispersed throughout the body. However, the education and activation of these immune cells are closely controlled by complex spatially organized tissues. One striking example is the role of secondary lymphoid organs (SLO) as pivotal sites for initiating immune responses. These specialized organs, which include lymph nodes, the spleen, and Peyer’s patches, gather antigens and cells from nearby tissues, recruit B and T cells from the bloodstream, and create an environment conducive to interactions between antigen-presenting cells and lymphocytes, ultimately facilitating the growth and specialization of effector cells.

This function is permitted by a particular architecture common to all SLO with distinct B cell follicles and T cell zones, supported by follicular dendritic cells and fibroblastic reticular cells respectively. Within SLO, antigen-presenting cells, like dendritic cells, strategically position themselves to efficiently present antigens to lymphocytes. High endothelial venules, specialized blood vessels that enable the entry and exit of lymphocytes, facilitate the trafficking of immune cells in and out of these organized structures (***2***).

While SLO develop at predetermined locations during embryogenesis, tertiary lymphoid structures (TLS) arise at sites of persistent inflammation induced by triggers such as chronic infection, malignant tumors, and autoimmune diseases (***3***). Although TLS can be found at various levels of maturity, at their most mature state they display T-cell-rich areas, where T-cells are clustered with mature dendritic cells, and B-cell-rich areas where proliferating B cells are found in a network of follicular dendritic cells, as well as high endothelial venules, making them structurally very similar to SLO (***4***). This, as well as their association with a mostly favorable prognosis in cancers (***5***) and an unfavorable one in autoimmune diseases, has led to the characterization of TLS as local drivers of antigen-specific immune responses in the tissue. Presence of TLS, their maturation stage, and cellular composition were all linked with positive clinical outcome and show high potential as predictive biomarkers (***6***). This thus makes them a high-priority target for translational research, as the modulation of the development of these structures could potentially be used to enhance or lessen antigen-specific responses in patients.

To gain a deeper understanding of these spatially organized structures, it is critical to employ not only single-cell methods but also spatially resolved approaches (***6***). With these innovative methods, it is possible to delve into the formation and maturation of TLS with unprecedented detail. Comparing gene expression profiles across TLS of different maturity levels, as well as contrasting them with the well-established SLO, offers a unique opportunity to decipher the intricacies of these organized immune microenvironments. This holds significant clinical implications, as understanding the molecular cues that lead to the development of TLS could lead to new therapeutic strategies aimed at enhancing the immune response against cancer by inducing TLS formation and maturation. Such experiments could also shed light on the cellular players involved in orchestrating these structures and how they communicate. Expanding into spatial transcriptomics, with its increased number of targets, high sensitivity, and spatial resolution, has the potential to deepen our delicate interplay between TLS, cancer progression, and therapeutic outcome.

### Single cell technologies and the definition of cell types and states

The existing landscape of spatial technologies and data analysis is already summarized in many excellent reviews (***7–12***), also with a focus on tumor microenvironments and TLS (***13, 14***). Here we will focus on aspects relevant to this chapter, and thus we will not provide a comprehensive list of references, and we would like to refer the interested reader to the cited references above.

Cells are the basic structural and functional units of organisms and recent breakthroughs in single-cell technologies have revolutionized our understanding of cellular function. Cells are commonly classified into different types, where each type refers to a group of cells performing similar functions that are distinct from other types. While being an incredibly useful description to investigate complex multicellular organisms, there is no consistent and standard definition of cell types (***15–17***). Single cell approaches now enable the characterization not only of the molecular composition of individual cells but also of their spatial context with increasing detail, yielding an increasingly nuanced description of their current cellular state. The ensemble of probed cells thus reveals a vast continuum of cell states, and novel methods allow the inference transitions between these states. The challenge now is to link these states with a robust and reproducible definition of cell types. In such a definition, a cell type could be considered generally stable and maintained by a set of cell-type-specific transcription factors. These cell types can exhibit different states associated with transient phenomena such as replication, differentiation or migration, responses to environmental cues, as well as processes related to development or disease. The investigation of these different states of cell types could allow the differentiation from core regulatory factors maintaining these cell types from factors driving specific functional states (***15***). By borrowing concepts from taxonomy, cell types can then be described in a hierarchical manner, progressing from broader classes to more specific subclasses with an increasingly fine-grained division.

### Spatial RNA profiling

Spatial RNA profiling not only provides the expression profile of individual cells, but also their spatial location. This hence allows not only to investigate the cellular heterogeneity, but also the spatial organization of these cells and their neighborhood relationships. Such data therefore allow to establish cellular architecture of embryos, tissues, or tumors and are crucial to understand tissue function (***1, 14***). Spatial RNA profiling can be divided into two types of approaches: first, untargeted sequencing-based approaches, and second, targeted hybridization-based approaches. The former have the advantage of providing a full transcriptomic view, but with reduced sensitivity and spatial resolution, while the latter are panel-based approaches allowing the detection of a predefined set of RNA species, but with higher detection efficiency. Commercial systems now exist for both types of approaches, providing turn-key solutions. However, these systems are often expensive, both in terms of instrumentation and reagents for experiments. Exploratory analysis, time-courses, dose-response, and replicates thus require substantial financial resources. Further, the counterpart of being relatively easy to use is a lack of flexibility, both for microscopy, e.g. the imaging modality, and the experimental protocol, e.g. to add signal amplification steps. Lastly, these approaches provide increasingly large panels, which are often required, but not all studies require such depth: smaller but more flexible panels are sometimes more desirable. For these reasons, we argue that home-built solutions, both in terms of instrumentation and experimental workflows remain an interesting choice for many research questions, particularly given the ever-increasing panel of exciting open-source analysis tools that are available. While such DIY approaches are more complex to set up, they are cheaper and more flexible, and will also help to democratize access to this technology.

In this chapter, we hence present a do-it-yourself (DIY) guide for spatial and single-cell RNA profiling. Our goal is to lower the initial threshold to get started by providing a step-by-step guide with recommendations. We will use TLS as a case example to illustrate the kind of questions that can be answered, but this approach can equally be applied to other biological models with complex multicellular structures.

### DIY spatial RNA profiling: overview

Spatial RNA profiling allows the localization of individual RNA molecules of different marker genes in native biological samples, such as tissue or embryos, with high spatial accuracy. These marker genes are either already known, or are inferred from scRNA-seq. These data can then be used for the identification of different cell types and their states in heterogeneous biological samples (**Figure 1A**). In this section, we will provide a summary of the entire workflow establishing the key concepts and terminology, each step of the workflow is then described in more detail in the following sections, pointing the reader to relevant literature and methods.

**Figure 1.**
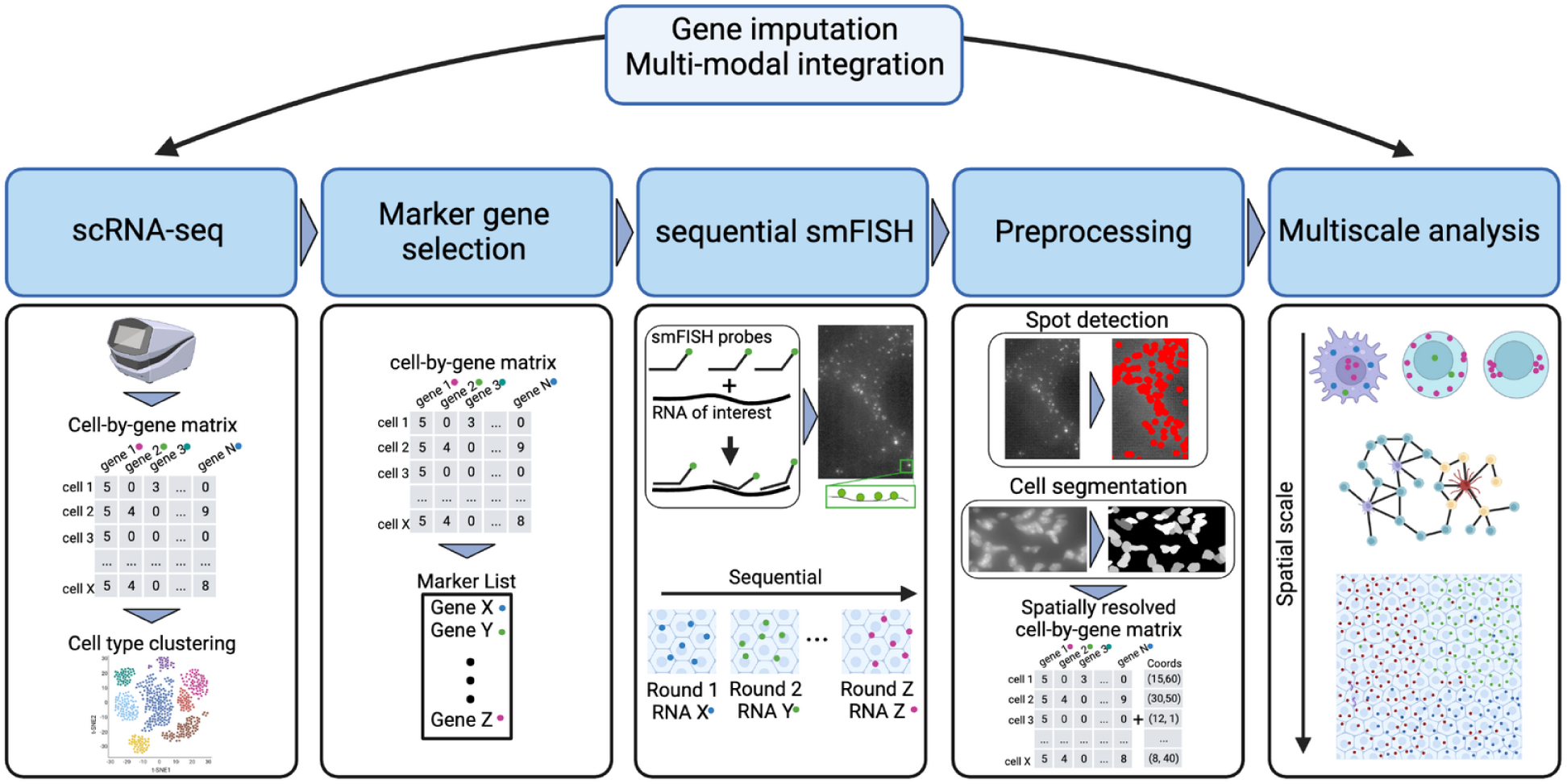
Summary of the spatial RNA profiling workflow presented in this chapter. Each step is detailed in a corresponding section.

The first important step is to determine the marker genes that allow the identification of the cell-types/states of interest. This can either be based on a priori knowledge or by analyzing other available resources, such as scRNA-seq data. The latter permits to establish a combinatorial list of markers with an optimized predictive power.

The RNAs of these marker genes are then visualized with variants of single molecule FISH (smFISH). For each target RNA, several short DNA oligonucleotides (probes) are bioinformatically designed and can be fluorescently labeled with different strategies. Using a sufficiently large number of probes gives strong local signal enrichment compared to individual non-specifically bound stray probes or autofluorescence. Samples can be imaged under a conventional fluorescence microscope, and individual RNAs appear as bright diffraction-limited spots.

On conventional microscopes, only a few (typically 2-4) fluorophores can be spectrally differentiated and imaged at the same time. However, the number of marker genes is often substantially larger. To image tens to hundreds of RNA species, iterative approaches have been developed alternating between probe-hybridization against a selected number of RNAs, imaging, and removing the previously used labels. In the simplest implementation, in each hybridization round the number of targeted genes corresponds to the number of available fluorophores (iterative FISH). The total number of targeted genes therefore scales linearly with the number of imaging rounds. To substantially increase the number of genes, multiplexed approaches have been developed where each RNA can be detected multiple times (typically 4 times). For each RNA a unique readout signature among the different hybridization rounds is then defined. In each hybridization round, a defined subsets of RNAs are detected. Post-processing then reveals the molecular identities of each identified spot. By using error correction methods, hundreds of distinct RNA species can then be detected in typically fewer than 20 rounds. The choice between iterative and multiplexed FISH depends on the targeted genes. Iterative FISH is suited for studies with not more than a few tens of target genes. It has the advantage of reduced experimental complexity and is less sensitive to highly expressed genes. Multiplexed FISH opens the door to studying hundreds to thousands of genes in the same cell and thus provides a more comprehensive view of the transcriptome.

While such experiments could be performed manually, they are often automated by using a microscope equipped with a computer-controlled fluidics system. These fluidics systems perform all buffer exchange, and when combined with adequate control software, enable a complete automation of the hybridization-imaging cycles.

The acquired data sets are large and are ideally analyzed with automated software tools with only minimal user interaction. While there are several methods available, the workflow of this analysis is relatively standard. In the initial analysis, the positions of RNA molecules are identified and can be treated as point clouds and statistically analyzed. Often, additional cellular markers such as a DAPI or membrane stain are added. Accurate cell segmentation is often challenging, and analysis methods are required to identify the expression profile and spatial location of individual cells. Once this is established, these data can be quantitatively analyzed with a vast range of methods, often based on advanced statistics and machine learning. Here, the choice of methods strongly depends on the nature of the data, e.g. the number of marker genes, the quality of cell segmentation, the spatial resolution, and the scientific question at hand, such as cell-to-cell communications, cell-composition, or large-scale tissue gradients.

## DIY spatial RNA profiling: a step-by-step guide

In the following sections we will provide detailed explanations for the steps described in the overview above.

### Marker gene selection

The RNA profiling method described here is a targeted approach against a selected ensemble of marker genes. The choice of these markers has to balance two important aspects: it should contain enough markers to permit a statistically robust identification of the cell types/states of interest, while being short to reduce experimental cost and complexity. These markers can be already established and validated genes, e.g. as signature genes for TLS (***5, 18***) or subsets of B-cells within human tumors (***6***). Alternatively, computational approaches can be used to identify new potential markers from other experimental data (such as scRNA-seq). In general, these methods can be divided in two classes: supervised methods based on predefined cell types/states, and unsupervised methods that establish the optimal set of cell types/states based on the input data. In the following paragraphs, we will describe their respective strengths and limitations and propose an approach to quantitatively compare these methods on simulated data, providing guidance for the final choice of markers.

#### Supervised methods

These approaches rely on prior annotations of the cell-types, which are typically obtained from scRNA-seq data. They then use different statistical approaches to identify a minimal set of marker genes that allows the identification of the annotated cell clusters.

A widely used approach is the identification of **differentially expressed genes (DEG)**, e.g. as implemented in popular tools like Scanpy (***19***) and Seurat (***20***) and benchmarked here (***21***). DEG methods are based on statistical tests to detect genes with distinct distributions in two cell populations. For an application in spatial RNA profiling, these approaches have several limitations. Most importantly, the user is required to manually choose the number of genes or to manually fix thresholds on computed statistics. Moreover, each cell cluster is treated separately, and thus no optimal combinatorial set of markers to discriminate all clusters is obtained.

To partially resolve these limitations, another type of supervised approach involves training a **machine learning model to predict the annotated clusters** from the gene expression matrix (***22–24***). Then, the most informative genes for this classification can be selected as markers, providing a solution to the computational complexity. One of these methods, NS-forest (***24***), is well adapted for smFISH experiments since it identifies genes that are not only information-rich but also prioritizes genes with binary expression which are more robust in the subsequent analytical tasks. An intriguing alternative is to leverage the hierarchical order within cell type annotations, e.g. sub-cell types, which could improve classification accuracy while reducing the number of required marker genes, as done by scGeneFit (***25***) or gpsFISH (***26***)

#### Unsupervised method

The previously discussed approaches have several limitations. First, they are not well suited to identify markers for rare molecular profiles. Cells with these rare profiles might not form distinct clusters in the high-dimensional gene space, leading to the absence of annotations. Secondly, supervised methods select optimized markers for prior annotations. However, they are less inclined to uncover genes that characterize subtypes or states across the classes as these might not be annotated, e.g. senescence or proliferation.

Unsupervised marker gene selection can address some of these challenges. Their underlying principle is to identify markers that summarize the full diversity of the transcriptomic profile in an input scRNA-seq data set (***27, 28***). As an example, geneBASIS (***28***) generates a k-nearest neighbor (k-NN) graph with the full set of genes as coordinates and selects a minimal set of markers that span a similar k-NN graph.

#### Challenges due to differences in experimental modalities

Most of the methods discussed so far do not consider the shift in experimental modality between single-cell RNA sequencing and imaging-based RNA profiling. First, most imaging-based methods do not cover the entire transcriptome in contrast to scRNA-seq. Second, the detection sensitivity of both methods is different, with imaging-based methods being more sensitive. Some recent methods start to address these challenges. PERSIST (***29***) employs a deep neural network predicting the full transcriptome by taking only a few genes as input. The gene list found to predict the full transcriptome is selected as the list of marker genes. To ensure the selected markers are effective in a spatial transcriptomic context, PERSIST first binarizes gene expression by considering genes to be either silent or expressed to address the domain shift from scRNA-seq to FISH measurements. It then incorporates a loss function that accommodates noisy gene dropouts in scRNA-seq data. Another method, gpsFISH (***26***) assesses possible gene expression discrepancies in targeted RNA profiling data compared to scRNA-seq and selects gene panels that are robust against predicted platform distortion in terms of sequencing depth at the gene level. To do so, gpsFISH compares scRNA-seq with existing image-based RNA profiling datasets and models the multiplicative and additive platform effect of each gene. Lastly, gpsFISH also selects genes in hierarchical order to address the above-mentioned challenge of sub-cell types.

#### Optimized markers for downstream analysis

With the exception of NS-forest (***24***), PERSIST (***29***), and gpsFISH (***26***), most methods primarily select markers based on their informativeness without considering whether these markers would facilitate downstream task analysis or account for the modality shift between scRNA-seq and spatial RNA profiling. Specifically, one of the primary challenges in RNA profiling lies in accurately assigning detected RNA molecules to individual cells (***30***). This task is complicated by the complexity of cell boundary staining, which is often not available, making RNA-cell assignment a non-trivial problem. In the absence of cytoplasmic staining, several methods have been explored and detailed in the subsequent section for single-cell RNA profiling. However, these methods are susceptible to errors influenced by both RNA expression patterns and cell morphologies. Such inaccuracies in RNA profiling potentially impede downstream analyses, particularly in tasks related to cell type/state identification, as noted in the literature (***31***)

#### Using simulation to evaluate the expected performance marker genes

One major challenge for image-based data, as detailed further below, is to correctly assign RNAs to their respective cells. Accurate RNA-to-cell assignment is crucial since this directly impacts the gene expression profile of each cell and thus all downstream data interpretation. As detailed in the dedicated section below, this assignment can be performed by using the positions of the RNA molecules and is thus influenced by the available fluorescent stains, gene markers, and tissue morphology. In this context, it is essential to estimate the extent to which potential errors in RNA-to-cell assignment can affect downstream analysis tasks, such as cell type/state identification. We believe that in-silico simulation can help to answer this question. For example, simulations of cells and their RNA transcripts were used to test if the selected marker could effectively identify the intended cell type using only nuclear staining (***32***). Similarly, simulations can help select the minimum number of markers (***33***). To better illustrate this approach, we provide along with this DIY guide a workflow as Python Jupyter notebooks, to simulate a smFISH experiment with a user-provided gene list **(Figure 2A**, https://github.com/tdefa/SimTissue). Simulated data can then be further analyzed to assess the expected performance obtained with the chosen gene marker list and RNA-cell assignment method (**Figure 2B-C**).

**Figure 2.**
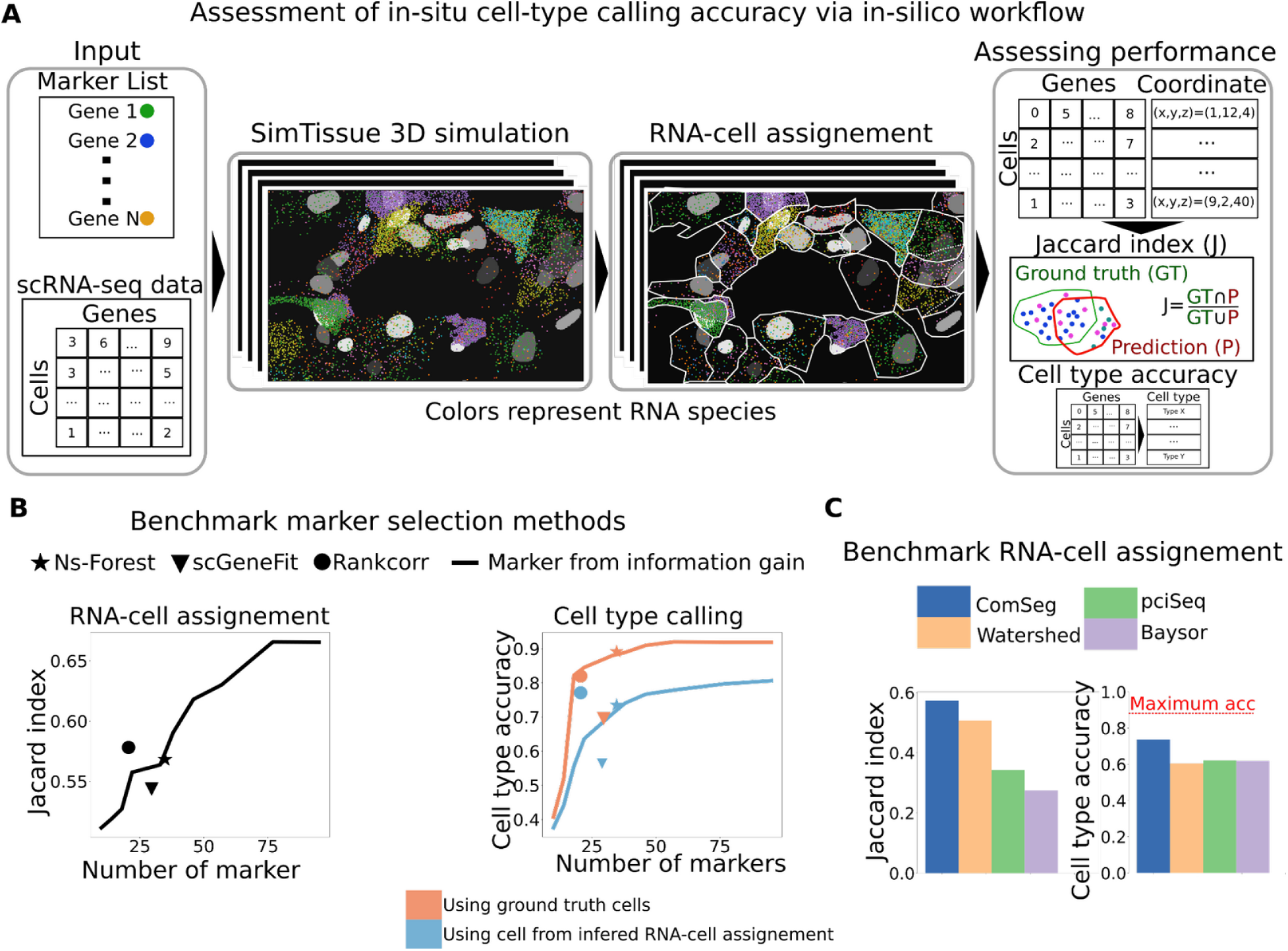
Using simulations to benchmark marker genes and cell segmentation methods. **A** List of user-provided marker genes is used to simulate smFISH images. Optionally, realistic RNA levels are drawn from scRNA-seq data. Simulation framework provides cell volumes and nuclei. RNA-to-cell assignment can then be compared quantitatively to the known ground truth with metrics such as Jaccard index. **B** Simulation of 19 cell types as identified from our recent study (**34**) in mouse lung tissue. Cell-type and Jaccard index (left) cell type accuracy (right) are shown as a function of the number of marker genes selected based on their information gain (IG, more details on GitHub). Orange curve assumes perfect assignment, blue curve obtained with our method ComSeg (**31**). Comparisons of three different marker gene selection methods (Ns-forest - squares, RankCorr - circles, scGeneFit - stars) are color-coded with either ground truth or analysis results. Results indicate large differences in the number of markers and their performance, helping to establish marker lists suited for imaging experiments. **C** Simulation workflow can also be used to benchmark different cell segmentation methods (ComSeg, PciSeq, Baysor, and Watershed). Results for simulations as in B, with marker genes obtained from NS-forest.

### Sequential smFISH experiments

These experiments permit visualization of the identified marker genes in the biological sample. If the number of markers is larger than the number of spectrally resolvable fluorophores, sequential approaches are used (**Figure 3A**) relying on alternating cycles of probe hybridization, imaging, and probe removal.

**Figure 3.**
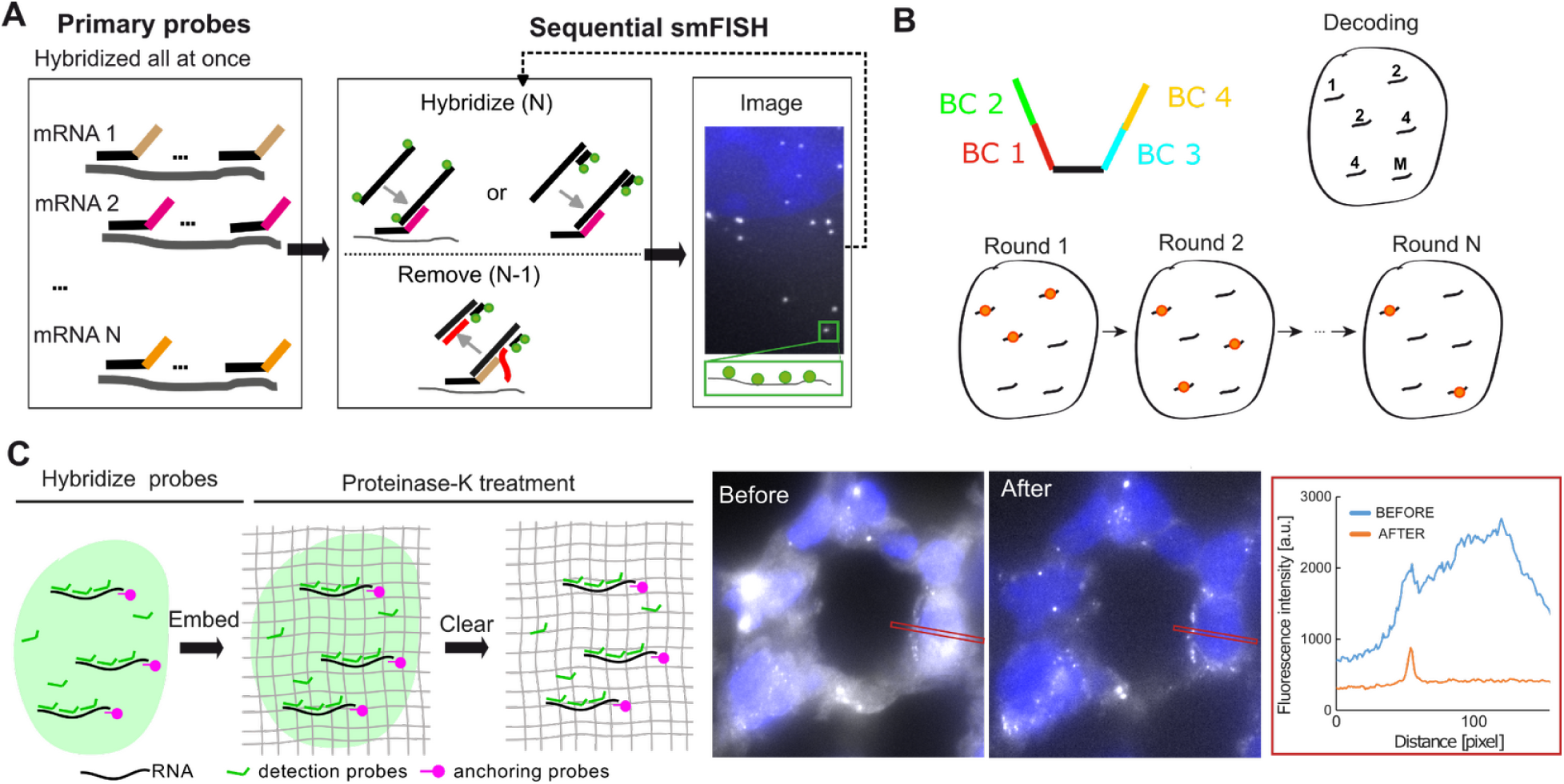
Sequential smFISH experiments. **(A)** Overview of typical experimental workflow. First, all primary probes targeting the marker genes are hybridized. All probes targeting the same gene have the same barcode (colored bars). In round readout probes for specific markers are hybridized, imaged, and stripped. **(B)** For multiplexed FISH, each RNA species has a unique combination of barcodes (typically 4). Several different RNA species are then visualized in each round, and the molecular identity of each spot is determined with an analytical decoding step. (**C**) (left) For optical clearing, tissue is embedded in a polymer matrix and RNAs are anchored. Sample is treated with proteinase K for several hours. (right) Effect of tissue clearing on smFISH signal. Selected z-slice of tissue before (left) and after (right) is shown. Note that different intensity scaling was applied. Rectangle indicates the region where intensity profile was measured. This profile shows drastic reduction in background and signal of individual RNA is clearly visible.

#### smFISH probes - design and synthesis

The RNAs of each identified marker gene are then targeted by several bioinformatically designed oligos (**Figure 3A**). Several different tools are available to design these probes that optimize probe sequences based on different criteria, such as GC content and melting temperature, often combined with additional bioinformatic analysis, e.g angler-FISH (***35***), OligoStan (***36***), PAINTSHOP (***37***), ProbeAngler, or ProbeDealer (***38***). While the users have a large choice, future efforts for a careful comparison of these methods and the establishment of a gold standard for probe design could further help to increase the sensitivity of these approaches.

These primary oligos can then be purchased from different commercial vendors. Depending on the length and the number of probes, different synthesis possibilities are available. Probes can be ordered on **96 well plates** with one or two oligos per well. While this usually gives sufficient probes for several thousand experiments, this is only suitable for relatively short oligos (<60 nts) and these plates are rather cumbersome for storage. A convenient alternative are **oligopools (opool)**. Here, substantially longer probes can be synthesized, and are provided as a mixed pool in one tube. The amount of provided probes depends on the synthesis scale. For smaller-scale pools with only a few hundred probes, a production scale of 50 pmol per probe allows to directly use the opool for smFISH experiments. Further, ordering one opool per gene is often not much more expensive, and permits to use pools individually, e.g. for validation, or creating custom pools for selected genes. For larger-scale studies with several thousand oligos, larger opools with lower concentrations can be purchased. These then require amplification before using them in smFISH experiments (***39, 40***).

#### smFISH probes - fluorescent labeling

The primary probes from above are not fluorescently labeled but carry one or several overhang sequences (termed "barcodes"). For iterative FISH, these barcodes are gene-specific and are the same for all oligos targeting the same gene (**Figure 3A**). For multiplexed FISH, each gene carries multiple barcodes which allow targeting the gene several times during the different hybridization rounds (**Figure 3B**).

To create a fluorescent signal, fluorescently labeled readout probes are used. These readouts can be either directly labeled with a fluorophore or carry an additional sequence that can be recognized by a third imaging probe (**Figure 3A**). Using a labeled barcode is simpler, with potential advantages in reduced background in tissue but comes with an increased price tag since labeled oligos are more expensive. Using a labeled imager reduces cost since the same imager can be used for different barcodes. It also makes it easier to change fluorophores since only a different imager probe has to be purchased.

#### smFISH in challenging samples

With the above-described detection strategies multiple probes per RNA are used, with each probe labeled with a few fluorophores. However, this signal might be too weak for detection, especially in more challenging samples such as tissues. Several approaches exist to circumvent this problem by either increasing the specificity of the primary probes, amplification of the signal, optical clearing of the sample, or a combination of several of these aspects (***41***).

Detection of the signal of individual RNAs can not only be hampered by autofluorescence of the specimen but also by signal stemming from non-specifically bound primary probes. Such non-specific signals can be reduced by utilizing paired primary probes (***42, 43***). Only if both probes are present, a bridge sequence can stably hybridize which is then decorated by fluorescently labeled probes. Several approaches for smFISH signal amplification were developed, which differ in experimental complexity, cost, and suitability for sequential FISH experiments, including complex protocols combining split primary probes with massive signal amplification such as clampFISH (***44***) and HCR (***45***). Other methods provide more moderate amplification, but can be more readily integrated in an already established standard smFISH workflow, such as branched DNA amplification (***46***) or SABER (***47***).

Lastly, tissue can be cleared, and proteins and lipids removed (**Figure 3C**). This not only reduces autofluorescence but also reduces non-specific binding of smFISH probes to these structures (***48***). These clearing approaches are based on the same underlying principle where RNAs are anchored to a polymer matrix. This anchoring can be achieved with chemical compounds (***49, 50***) or anchoring probes targeting the Poly-A tails of mRNAs (***48***). Once embedded the specimen can then be exposed to rather harsh treatments involving high concentrations of SDS and ProteinaseK. This effectively removes lipids and proteins from the sample, while the anchored RNAs remain in the polymer (**Figure 3C**).

#### Signal removal

To remove signals of the preceding hybridization round, several approaches are commonly used: high-stringency wash buffers, chemically cleaving, or probe-stripping. The advantage of these approaches over photobleaching is that the entire sample is treated at once and no specific light source is required.

High-stringency buffers, usually with high formamide concentrations, can be used to remove probes. Depending on the experimental design, either only the barcode probes (***51***), or all probes, including the primary probes (***52***) are removed. In the latter case, primary probes for the next round need to be hybridized resulting in longer experimental workflows. Alternative approaches relying on less toxic agents and fast removal are available.

In chemical cleaving, the fluorophores are conjugated to the probes with a disulfide linkage. Interestingly, the fluorophores can then be rapidly cleaved from these probes with a short Tris(2-carboxyethyl)phosphine (TCEP) incubation (***53***). While these oligos are more expensive, cleavage is rapid and complete since the cleaved oligos can be easily washed out.

In probe-stripping, probes (typically the readout) contain an additional sequence termed "toe hold". By hybridizing stripping probes that are complementary to this toe-hold and the barcode sequence on the primary probe (**Figure 3A**), the readout oligo can be removed (***40***). Since stripping of the previous round and hybridization of the next round can be performed simultaneously no additional steps are acquired. This approach is simple and inexpensive due to the low cost of the unlabeled stripping probes.

#### Experiments: bench and automation

In the actual smFISH experiment, samples are fixed and permeabilized (***36***). Then all primary oligos targeting the genes of interest are hybridized. This hybridization is performed on the bench and can take from several hours to several days. After washing, the samples can then be exposed to the subsequent rounds of hybridization, most frequently by using a computer-controlled automated system combining a microscope and a fluidics system. These systems permit automated buffer exchanges on the microscope, and samples are typically placed in a closed heated imaging chamber creating a reaction volume of a few hundred uL (***54, 55***) Fluidics systems can be either commercial (***55***) or custom-built (***52, 54, 56, 57***). For the latter, often detailed building plans and control software are available. Communication with the microscope can be achieved in different ways: either by providing a fully integrated solution including microscope control (***54, 56***), using open-source acquisition software such as PycroManager (***57, 58***), or providing different options to trigger acquisition in microscopy control software such as TTL or synchronization files (***57***). The latter has the advantage that they can be easily adapted to any microscope that provides the possibility to trigger acquisition with an incoming signal. In summary, these fluidics systems can be combined with already available microscopes, and thus provide a cheap but also flexible solution, since the imaging modality, e.g. wide-field, spinning disc, or light-sheet, can be changed. Importantly, the user has full control over the image acquisition, e.g. to also record imaging stacks of thicker specimens.

### Preprocessing: identify single-cell RNA expression profiles

After image acquisition, cells and their expression profiles have to be identified. This usually encompasses detection of RNA molecules, the segmentation of cellular landmarks, such as nuclei and the assignment of RNAs to their respective cells (**Figure 4**).

**Figure 4.**
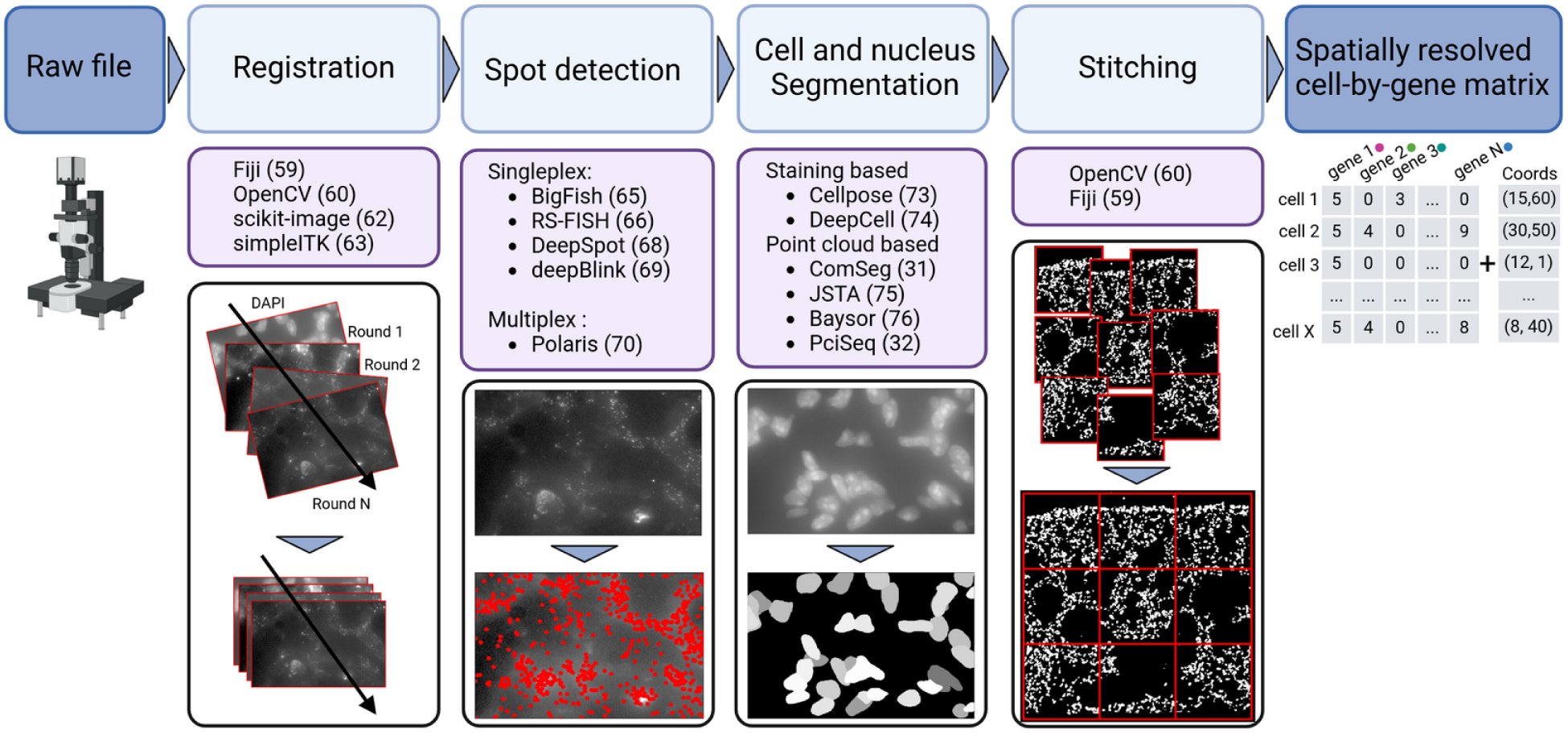
Preprocessing of spatial RNA imaging data. These workflows use raw image files as input and provide, as a final result, a spatially resolved gene expression matrix for each cell.

#### Image stitching and registration

Acquisition of large fields of view requires stitching of the individual tiles. If stitching is not automatically performed by the acquisition software, different open-source tools exist, e.g. in Fiji (***59***) or OpenCV (***60***). Further, images acquired for different hybridization rounds need often to be registered due to drift of the microscope stage. Such registration can be performed on existing landmarks in the images, e.g. nuclei, or if high precision is required by adding dedicated fiducial markers, i.e. bright fluorescent beads. Similarly, several methods for image registration are available in Fiji (***61***), scikit-image in Python (***62***), or frameworks such as openCV (***60***) and simpleITK (***63***) with programming interfaces in many languages.

#### RNA detection and decoding

In smFISH images, individual RNA molecules appear as diffraction-limited spots. Spot detection has been extensively studied in image processing (***64***) and several dedicated analysis packages for smFISH are available. For instance, our package FISH-quant (***65***) utilizes a well-established approach for spot detection, implemented in Python, and provides documented and modular end-to-end analysis workflows. RS-FISH (***66***) is implemented as a FIJI plugin (***67***), optimized for fast detection and distributed computing. One major drawback of these similar approaches is the need for human intervention to set or verify detection thresholds. More recently, several deep-learning-based approaches have been presented (***68–70***). These methods are trained on increasingly larger annotated data (both simulated and experimental) and should thus in principle be directly applicable to new data. Some approaches provide web-based user-interfaces for rapid testing, without the need for installation. These approaches will certainly be widely used, retraining on new data and efficient extensions to 3D will be important aspects. A particular challenge for these data is the creation of an experimental ground-truth since manual annotation of thousands of smFISH spots per image is not realistic. Consensus annotation across several analysis packages provides an intriguing, automated solution (***70***). Lastly, providing an easy-to-use interface for retraining will permit end-users to further adapt these approaches to their data if required. Lastly, if RNA species are multiplexed, a gene inference step is required to infer the molecular identity of each RNA spot after detection in the multiple hybridization rounds.

#### Nuclei segmentation

Deep learning-based approaches are most widely used for nuclei segmentation, and provide accurate segmentation with pre-trained models, even on 3D data (***71–74***). Recent implementations now also permit rapid retraining on small, annotated data sets further optimizing the performance for specific data sets (***73***).

#### Cell segmentation

The previous steps provided nuclei segmentation and the positions of RNA spots. The last crucial steps associate RNA molecules with cells. The correct estimation of cellular RNA expression profile is crucial, as errors in this assignment will hamper all other downstream analyses. A first solution is to use cytoplasm boundary staining and use deep-learning models for cell segmentation (***73, 74***). However, such staining is not always consistent among different cell types, and certain cell morphologies in tissue can be very challenging to segment (***74***). Therefore, new methods have been presented that perform cell segmentation directly on the RNA points clouds. For instance, PciSeq (***32***) and JSTA (***75***) identify cells with similar expression profil as in annotated clusters from scRNA-seq data. Further, PciSeq imposes a circular shape prior. In contrast, Baysor (***76***) and ComSeg (***31***) do not rely on external datasets. Baysor finds cells with elliptical shapes that have homogenous expression profiles by using maximum likelihood estimation (i.e. using statistical modeling of a cell). ComSeg assumes no prior shape and associates RNAs to nuclei with a high estimated probability of being co-expressed in a given cell. Of note, simulations can help to benchmark different cell segmentation methods (**Figure 2**).

#### Data normalization

As for scRNA-seq data, the estimated cell expression vectors after segmentation may contain experimental artifacts. In the context of smFISH, artifacts may come from variation in the RNA probe sensitivity, missed detection of RNA molecules, errors in the RNA-to-cell assignment or batch effect across different tissue samples. To remove potential gene variance from experimental artifacts and keep only gene variances that are biologically meaningful, normalization methods can be employed. Most of these methods were developed for scRNA-seq data (***77***) but can be also applied to spatially resolve single cell RNA profiles. For example, the scTransform (***78***) has been successfully used to classify spatially resolved RNA profiles (***79***) in imaged-based data.

However, methods for scRNA-seq can assume that the entire transcriptome is covered, and total RNA count per cell as a proxy for sequencing depth to then infer scaling factors. However, direct application of these methods on imaging-based data can be problematic due to their limited coverage of the transcriptome. Importantly, application of scRNA-seq normalization methods on imaging data can lead to erroneous biological conclusions (***80***). An interesting future avenue could be to consider morphological information of the cells. For instance, recent studies illustrate that RNA counts for many genes are strongly correlated with cell volume (***81***). As an example, stLearn through stSME (*S*patial *M*orphological gene *E*xpression) proposes a dedicated normalization approach for spatial transcriptomic data by leveraging tissue morphology. stSME assumes that cells with similar morphology have similar RNA profiles. However, these morphological embeddings extracted with a convolutional neural network are obtained for small tissue sections and not single cells. Future iterations could estimate these features from individual cells.

### Downstream data interpretation

The preprocessing described in the previous section yields not only the gene expression matrix of individual cells but also their spatial position in the imaged sample. Further analysis then permits a quantitative description of the tissue architecture by leveraging both the molecular and cellular identity information as well as spatial descriptors. Importantly, these analysis approaches have to be carefully selected since they permit studying biological processes over different spatial scales (**Figure 5**), ranging from subcellular to cell neighborhoods to tissue gradients (***9, 14***). Lastly, this field is rapidly expanding and difficult to navigate for new users. Analysis frameworks such as Seurat (***20***) or Scverse (***82***) allow for a large range of analysis tasks thanks to shared data formats and community development guidelines.

**Figure 5.**
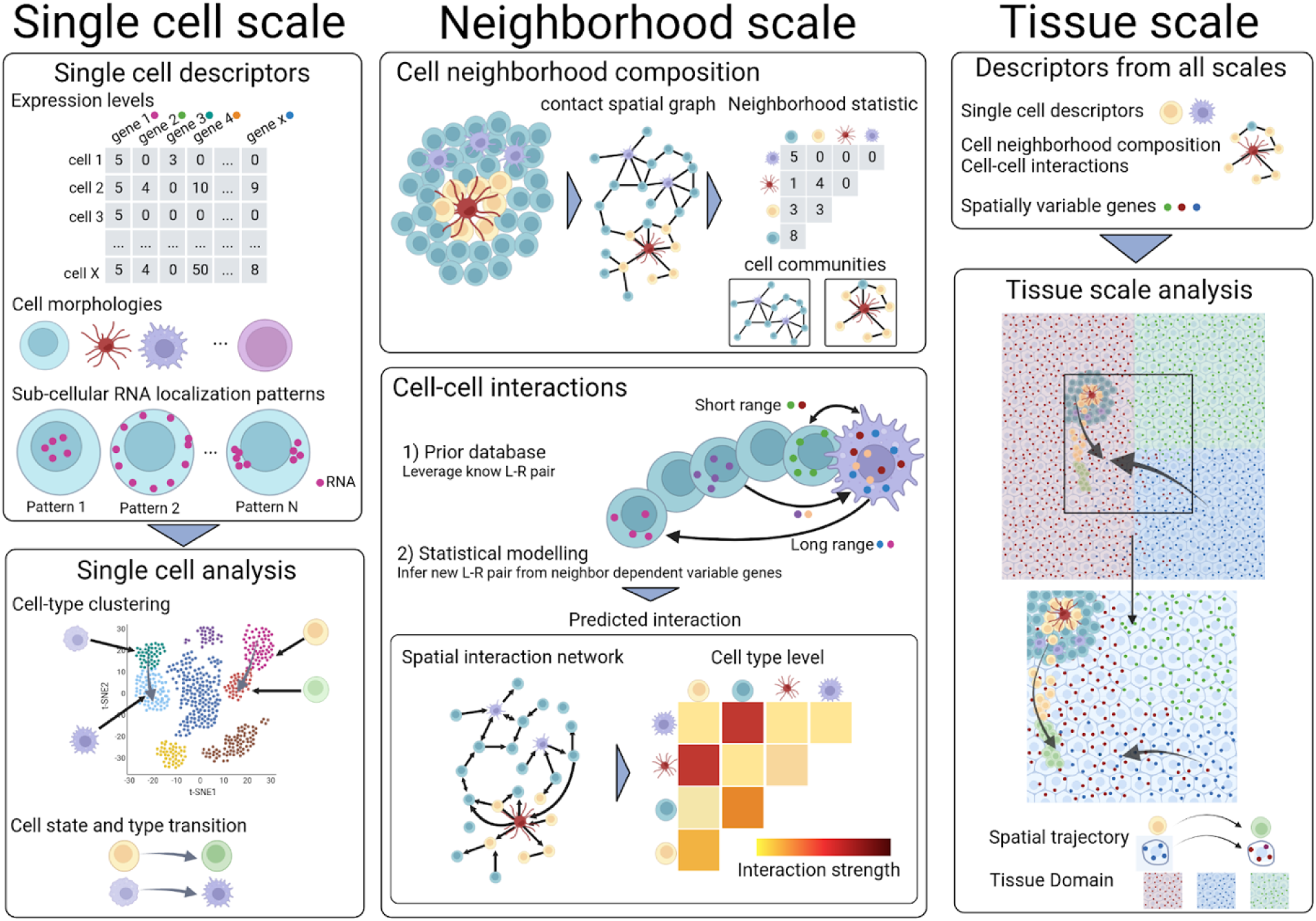
Downstream data analysis on several spatial scales. Each individual cell can be described by several quantitative descriptors summarizing the expression level of each RNA species, its cell morphologies and the subcellular localization of its RNA. These descriptors can be used to cluster cells and infer cell type and state lineage. The tissue can be described by a graph, where each node is a cell and neighboring cells are connected by an edge. Contact statistics among cell types can be computed and recurring patterns of cell communities be identified. Cell-cell interaction can be studied by combining existing Ligand-Receptor (L-R) databases or using statistical modeling. For the latter, the spatial position of cell types is analyzed along the cell expression level of each cell to identify neighbor dependent variable genes as candidate L-R. All the previously computed information can be combined with tissue scale descriptors such as spatially variable genes to analyze the entire tissue by identifying tissue domain and spatial trajectories.

The study of TLS and their interaction with the tumor microenvironment nicely illustrates why an analysis on different scales is important to obtain comprehensive biological insight. On the scale of individual cells, the functional state of B cells within TLS could be probed by designing adequate probes. By considering cellular environments, the cellular TLS composition and spatial reorganization in responses to therapeutic treatments such as immune checkpoint inhibitors can be studied. On even larger scales, TLS networks within a single tumor could be analyzed. Maintaining cellular resolution will be important as illustrated in a recent study of colorectal cancer (***83***), where it has been shown that the cellular composition of a TLS network within a single tumor can be heterogeneous.

### Cell type/state assignment with gene expression profiles

One of the most common analyses is to group cells in different clusters based on their expression profiles. However, there is no standard procedure to perform this analysis, which further depends on the number and identity of the marker genes (***14***).

This cell type assignment can be performed with methods also used in the analysis of scRNA-seq data (***84***). Most commonly unsupervised clustering of RNA expression profiles is performed with graph-based algorithms like Leiden (***85***) or Louvain (***86***). These methods have the advantage of adapting the number of clusters to the data without making assumptions about cluster size and shape. The clustering results can then be qualitatively assessed and visualized with dimensionality reduction methods like Umap or t-SNE (***87, 88***). Matching the clusters to known cell-types can either be done by manual annotation, or by using automated annotation tools. Numerous tools have been developed in the context of scRNA-seq for this purpose (***89***), such as scmap (***90***) or more recently scGPT (***91***), a foundational model for scRNA-seq trained on more than 10 million human cells. However, matching RNA profiles from smFISH and cell type annotations by using tools designed for scRNA-seq is not trivial. The main difficulty is the difference in sensitivity of both technologies and the number of measured genes. It is therefore advisable to use methods adapted to deal with the modality shift between scRNA-seq and spatial data like mfishtools (***92***) or the ones benchmarked here (***93***).

#### Integrating spatial information

The methods presented so far are based on the individual cellular expression profiles without considering the available spatial information, which can be leveraged at different scales as detailed next.

Spatial cellular context can be important, e.g. it is known that specific cell types reside together (***94***), or the activation state of (immune) cells can be better elucidated by considering their immediate neighborhood (***11***). An intriguing opportunity is to not only use the actual expression profile of cells to define their identity, but also include their neighborhood relationship with other cells in the tissue (***95***). Incorporating spatial features can help to obtain more robust cell type classification by reducing the contribution of expression noise, as shown by recent graph neural network approaches (***96, 97***). Further, spatial information can also help to define new sub-cell types with similar RNA profiles but distinct neighborhoods (***95***).

While the methods presented so far focus on a cell-level analysis, smFISH-based approaches provide data with high subcellular resolution. It is now well established that subcellular RNA localization is not random, but that RNA can localize dynamically to different subcellular compartments. Such preferential localization has been linked to transcriptional and translation regulation, local protein synthesis and complex assembly (***98***). Importantly, this localization can be dynamic, e.g. for development (***99***), and misregulation can cause disease (***100***). Importantly, it is possible to describe these subcellular RNA localization patterns with quantitative features allowing their integration in machine learning methods (***65***). Having simultaneous measurements of several genes also permits new analysis to identify subcellular regions with spatial proximity of genes, which can lead to improved cell classification (***101–103***). Interestingly, the identification of preferentially co-localized genes can yield testable hypotheses regarding their functional implications (***103***).

#### Cell state trajectories

Beyond cell type assignment, several approaches - mainly used in scRNAseq - can provide information about cell state trajectories. Their objective is to predict the future cell state or RNA profile and establish lineage relationships between distinct cell populations. Many methods like Monocle (***104***), Slingshot (***105***) model discrete cell state transitions to find lineage relationships in pseudotime. To achieve this, these methods construct a graph connecting the cell states, where the edge weights are estimated based on similarity of the gene expression profiles alone. This graph is then optimized to have minimum total edge weight and without any cycles resulting in the so-called minimum spanning tree. To our knowledge, the efficacy of these methods remains unknown when applied to image-based data characterized by a markedly reduced number of genes.

More recently, RNA velocity (***106***) uses the proportion of spliced and unspliced mRNA as an indicator of changes in transcriptional profile and models continuous-time transcriptome dynamics to predict the future state of cells. One limitation is its ability to predict future cell states only on the time scale of hours. Recent improvements extend the prediction time scale using ordinary linear equation (ODE) (***107***) or deep learning (***108***). While predominantly applied to scRNA-seq data, a recent study performed RNA velocity on imaging data, by leveraging changes of nuclear vs cytoplasmic as an indicator for changes in the transcriptional program (***109***). In the future, we believe that including changes in RNA localization as well as direct intron detection (***110***) holds great promise to further refine the definition of cell states and their transition.

Combining RNA expression profiles with spatial information at the scale of cell neighborhoods or the entire tissue allows to identify the spatial aspects of cell lineage relationship. Pseudo-time-space (PSTS) such as in the stLearn framework (***111***) methods can be used to describe spatial gradient of cell activation, differentiation or cancer evolution. Here, the edge weights in the graph are now not only calculated based on gene expression profiles but also consider the spatial distance between cells. Alternatively, spaTrack (***112***) employs an optimal transport framework, with the advantage of simultaneously analyzing multiple tissue samples, like pseudo-time tissue sample dataset. SpaTrack thereby enables the inference of a dynamic map of cell migration and differentiation across (pseudo)-time and tissue samples.

Of note, RNA velocity and cell lineage tracking are complementary in terms of relevant time-scales, as the first considers transcriptome dynamics in the range of hours, while the second is looking at longer time-scales with multiple cell type transitions

#### Spatial heterogeneity of cell types

An intriguing possibility to study tissue morphology is to describe the tissue as a spatial graph of cells. Such graphs can be efficiently built from 2D or 3D segmentation masks with packages as Tysserand (***113***) or Griottes (***114***). These graphs can then be used to identify spatial patterns based on cell-type heterogeneity (***115***). Alternatively, a community detection approach (***116***) can be employed to identify tissue regions exhibiting distinct patterns of cell type heterogeneity, subsequently linking them to specific biological functions. These approaches could prove particularly useful to explore TLS, since TLS were shown to be highly heterogeneous and can have different cell type composition within a single tumor and further exhibit intra cell-type spatial pattern (***83***).

#### Spatially variable genes

The comparison of the expression profiles between cells can yield highly variable genes (HVG) acting as a discriminative feature between different cell populations. A HVG is considered as a spatially variable gene (SVG) when it correlates with a spatial pattern. SVGs provide insight for the identification of tissue function or different cell-cell interaction patterns (***117***). One of the challenges of SVG methods is to separate the contribution of spatial variations from the intrinsic variation between cells. Early SVG approaches, such as SpatialDE (***118***), are based on hypothesis testing where non-SV is the null hypothesis and SV is the alternative. A *P*-value threshold is employed to control the false discovery rate (FDR). Recent methods provide more detailed information, for instance, CTSV (***119***) identifies cell-type specific spatially variable genes. SINFONIA (***120***) identifies SVG facilitating downstream task analysis like tissue domain identification (described in the next paragraph). A more extensive description of SVG methods can be found here (***121***).

Overall, we would advise to choose and employ SVG methods motivated by the underlying research questions. As an example, if the goal is solely to detect SVGs, we would advise to carefully control FDR rate while it is not necessary if the goal is to detect a low embedding of genes to perform downstream tasks like tissue domain identification or cell-cell communication (***120***). Besides, different SVG methods do not detect the same spatial pattern as shown in several benchmarks (***122, 123***). Applying several SVG methods and consolidating their results could hence provide a more comprehensive view of the different possible SVG patterns. Moreover, aggregating information from the SVG patterns can help for tissue domain identification as described in the next paragraph.

#### Tissue domains

Tissue domains are commonly defined as spatially continuous groups of cells sharing similar expression profiles. A range of methods permit to perform such spatial clustering. For instance, BayesSpace (***124***) and the Giotto (***125***), use a hidden-Markov random field (HMRF) to attribute a spatial domain label to cells based on their RNA profile. This attribution is determined by a strength parameter, which determines to which degree neighboring cells have the same domain membership despite differences in their expression profiles. Alternative methods, such as stLearn (***126***) or DeepST (***127***), include morphological cell features extracted with deep learning. These methods leverage the hypothesis that cells with similar morphology have similar RNA profiles. Morphological analysis can improve tissue domain identification as it enhances transcriptomic information, especially in tissue areas with poor RNA detection. In general, tissue domain analysis is thus best suited for samples with larger continuous areas of similar cells, and is less applicable for cases with very heterogeneous cell compositions in small neighborhoods, as it is the case for TLS (***83***).

#### Cell communication and regulatory pathways

Cell-cell interaction (CCI) can be defined as the changes in gene expression of a cell induced by the reception of a ligand from a sender cell. CCI can be inferred solely using the single cell RNA profile by leveraging prior knowledge from established databases of ligand-receptor pairs. However, considering the spatial position of the cell reduces the prediction of false positives by constraining the interacting cell to be close in space. For instance, Giotto (***128***) combines spatial proximity metrics between cell-types and ligand-receptor databases to better estimate communication between cell types. Similarly, COMMOT (***129***) inferred CCI in space with collective optimal transport enforcing limited spatial ranges and prior ligand-receptor pairs. Another approach, SpaGRN (***130***), also integrates prior knowledge on regulatory relationships and signaling paths to predict intracellular regulatory networks and identify their spatiotemporal variations.

While spatial RNA profiling can enhance the CCI prediction from existing ligand-receptor databases, it can also be used to estimate the length scale of interactions or even to identify genes involved in the CCI. SVCA (***131***) identifies potential genes potentially involved in interaction based on the assumption that the gene expression variation depends on intrinsic effects, environmental effects, and cell-cell interaction. SCVA identifies interacting genes as those with the expression variance being largely explained by cell interaction. Similarly, NCEM (***132***) leverages the same principle but permits setting the interaction scale as a parameter. CellNeighborEX (***133***) identifies gene variance due to adjacent cell type to uncover new genes potentially involved in cell-to-cell communication.

#### Multi-modal data integration

Increasingly, tissues are probed with different technologies to obtain a more holistic view of tissue function. Combining these data poses a formidable challenge for data analysis, and an increasing number of multi-modal data integration approaches have been developed (***134, 135***).

For imaging-based RNA profiling, an integration with scRNA-seq data can provide several advantages. First, this can improve and aid the cell type calling (***32***) or CCI (***136***). Second, this provides exciting opportunities for gene imputation, i.e. to predict the part of the transcriptome that has not been probed in the imaging experiment. SpaGe (***137***) projects the expression vector from imaging and sequencing data in the same space. It then uses the proximity of expression vectors from scRNA-seq and spatially resolved data to perform gene imputation. Alternatively, Tangram (***134***) associates each spatially scRNA-seq data to a spatial location by maximizing the gene spatial correlation between the chosen spatial transcriptomic dataset and the mapped cells from scRNA-seq. More recently CeLEry (***138***) is able to recover the location information of RNA profiles from scRNA-seq. CeLEry relies on ST datasets to train a neural network to associate sequenced cells with a spatial location and provides, importantly, uncertainty estimates for the recovered locations. Intriguing, multi-modal integration can also be used to combine technologies measuring different molecular aspects, such as immunofluorescence or chromatin accessibility (***135***).

### Analysis frameworks and data formats

We presented in the two previous sections a range of methods for the preprocessing and downstream analysis of spatial RNA profiling data. While the vast number of these approaches provides exciting opportunities, it can also be a daunting universe to navigate.

A user-friendly solution to this problem are frameworks regrouping different analysis tools. These frameworks use standardized data formats, and contribution guidelines to facilitate their usage. Of special note are Seurat and Scverse. Initially designed for scRNA-seq analysis, the popular R package Seurat has been extended to analyze spatial data (***20***). In addition to adapting its data format (*SeuratObject)* to store spatial data, it also offers spatial visualization and analysis like detection of SVG and integration with reference scRNA-seq to ease in situ cell type calling. Scverse is a broad ecosystem of packages, mainly written in Python, for single-cell omics data analysis (***82***). It builds on several core packages, such as Squidpy for spatial single-cell analysis, and different data formats for annotation (anndata (***139***)) and multimodal data (mudata). For new users, we recommend starting with these analysis frameworks, as they provide a large range of methods that can be selected according to the underlying biological question.

## Conclusion and perspectives

Spatial RNA profiling provides intriguing possibilities to quantitatively study tissue architecture and function. In this chapter, we present a comprehensive guide for how to perform and analyze such experiments with available open-source tools. This exciting field is continuously growing, and future developments, both experimental and analytical, will open exciting new possibilities and overcome current limitations. Beyond the potential perspective directly described in the sessions above, we believe several other directions are compelling.

Most studies and commercial systems perform 2D measurements, or probe only a thin 3D volume. However, observing thick 3D samples will be important in the future, e.g. to observe intact organoids or tumoroids. More specifically for TLS research obtaining large-volume images encompassing both TLS as well as the tumor microenvironment (TME) can be important. For instance, this would allow us to study the heterogeneity of individual TLS forming an interconnected 3D network within a single tumor (***83***). Further, spatial localization of TLS within the TME can have important implications in regard to anti-tumoral immune function, responses to immunotherapies and clinical outcome (***6***). Observing such thick samples poses challenges that can be addressed with recent developments. By combining adequate experimental protocols for tissue clearing and signal amplification with sensitive microscopy (spinning disc, or lightsheet microscopy) it is possible to observe single RNA molecules tens to hundreds of micrometers in tissue (***140***). However, imaging of such specimens will be slow. Increasing the number of channels that can be imaged in one round by combining fluorescence spectral and lifetime imaging promises to substantially decrease experimental time (***141***). These experiments create large image datasets requiring next-generation file formats such as OME-ZARR to address important issues of scalability (***142***). Lastly, the analysis tools need to be designed to handle such large data sets by allowing for distributed computing and GPU-acceleration for preprocessing tasks such as spot detection and segmentation and time-consuming calculations in the downstream analysis. Equally important will be tools for data exploration and interactive visualization.

Analysis of these increasingly large data sets will require the ongoing development of adequate tools, with the accurate integration of spatial information across scales (***9***). An interesting avenue to explore this large body of data could be inspired by the recent development in scRNA-seq, where large foundational models are developed like scGPT (***91***). Akin to large language models, scGPT is first trained on millions of cells with self-supervised pretext-tasks, such as predicting the entire transcriptome using only partial data, to learn a low dimensional representation of the data. Importantly, these tasks do not require annotations and can thus be run on extremely large data sets composed of millions of cells. Intriguingly, the model can then be fine-tuned without the requirement of large data annotation and outperformed the current state of this art for many downstream tasks like cell type calling or gene perturbation prediction. With the increasing number of datasets for spatial RNA profiling, we believe that foundational models like scGPT will be soon adapted for spatial resolved data with similar improvement for downstream analysis tasks.

The described method in this chapter focuses on RNA detection, but other published approaches combine FISH measurement of the transcriptome with simultaneous measurements such as epigenome, proteome, and metabolome. Analyzed with the adequate frameworks these data yield a multi-parametric and multi-scale description of cell-states at increasing fine graduation. Equally important will be the integration of dynamic information to access developmental relationships by following the lineage history of cells together with their molecular profiles (***16, 143***). Taken together, these new technologies will allow to establish a dynamic description of the cell states and possible trajectories therein. This promises to enable a refined understanding of the plastic nature of cellular function in healthy and diseased conditions.

We are in the middle of an exciting area with the advent of methods, both commercial and in-house developed, permitting to probe biology with high-spatial resolution across scales. However, the methodological landscape is becoming hard to navigate, especially for novices. Ongoing efforts with centralized frameworks such as Scverse (***82***) or Squidpy (***144***) are crucial since users can obtain guidance for methods choice and select different analysis methods without reformatting their data. The establishment of large consortia, such as the 4D nucleome program (***145***) or the Cell tracking challenge (***146***), could further help to organize developments and provide a community-driven exchange of validated protocols, building plans, and analysis tools. Lastly, it would also be interesting to organize benchmarking initiatives with data-analysis challenges that permit an objective comparison of different analysis approaches, akin to the Cell Tracking Challenge (***146***).

## Acknowledgements

This work has received financial support through the Agence Nationale de la Recherche (ANR) grants LUSTRA and TRANSFACT. F.M. acknowledges funding by Institut Pasteur. Figures 1, 4, 5 were created with BioRender.com.

